# Anticipating invasion of parasite with graph theory: *Dryocosmus kuriphilus*, a threat for *Castanea sativa*

**DOI:** 10.1101/2025.09.15.676224

**Authors:** Martin Rosalie, Jean-Loup Zitoun, Arnaud Casteigts, Sébastien Gourbière

## Abstract

The increase in world trade contributed to a rise in the number of insect biological invasions. The colonisation success of herbivorous pests depends mostly on the abundance of vegetal resource across the newly invaded territory. The explicit spatial representation of plant distribution offers a cost-effective approach to anticipate the spread of herbivorous insects over areas with fragmented and heterogeneous vegetal populations. In this article, we focus on the case of the recent invasion of chestnut trees (*Castanea sativa*) of the French Eastern Pyrenees by its most virulent pest, the gall-forming parasite *Dryocosmus kuriphilus*. Using tools from graph theory and available public data on the mesh size and distribution of chestnut trees, we model the spatial distribution of the pest resource across natural forests. In this framework, a graph provides a mapping of 1 km^2^ quadrats that constitute the territory. Quadrats are grouped to form communities, which are areas with homogeneous chestnut tree distribution, and in which the risk of parasite infestation and propagation is similar. Graph traversal algorithms then measure the vulnerability and the dangerousness of each patch, defined as their susceptibility to become infected and to contribute to the pest propagation. Such a spatial assessment of the invasion risk provides unique insights into potential propagation scenarios, improving the early detection and spread monitoring of these insect invaders.

**Author summary:** 

## Introduction

The frequency of biological invasions has risen steadily over the last two centuries, following the growth of international trade [1–3]. Invasive species are now widely recognized as a global threat, as they represent a leading cause of biodiversity loss in all natural ecosystems [4] and impose severe economic costs to agroecosystems (e.g. [5, 6]). While forests cover 30% of the world’s land area [7], providing 75% of the primary production and one of the largest carbon (C) sink on Earth [8], non-native species have become a significant part of the ‘mega-disturbances’ that impairs their health, ecological properties and associated ecosystem services [9]. Insects are the most species-rich group of forest invaders [10] and the number of species that have been transported away from their natural range has increased dramatically on all continents [1, 2, 4, 11–13], where they represent, along with invasive pathogens, one of the greatest threats to natural and managed forests [14, 15].

A critical challenge to improve invasion risk assessment and control strategies lies in the identification of the landscape features that can promote or impede the spatial spread of non-native species after the initial establishment of a local population [16–18]. The expansion of non-native herbivorous insects is restricted by the distribution of their host plant that constitutes a typical ‘bottom-up’ control of their population dynamics [19]. This provides a clear opportunity to integrate habitat suitability and connectivity to characterize the spatial network of host tree patches that underlie landscape susceptibility to insects’ invasion. Methods based on graph theory have received growing attention to understand the key anthropogenic and ecological determinants of biological invasions at various scales [20]. One of their most appealing application is in allowing to quantify connectivity between nodes representing habitat patches linked by arcs corresponding to potential dispersal paths [16]. This indeed offers efficient integrative modelling frameworks for identifying key habitat patches, facilitating early detection, and preventing secondary spread of an established invasive species [16], as well as prioritizing sites for control [21].

In this contribution, we aim to leverage graph theory methods to describe the network of habitat patches that has allowed the invasive chestnut tree insect pest, *Dryocosmus kuriphilus*, to spread through the natural forests of the French Eastern Pyrenees over the last decade [22]. This insect pest, a gall-forming hymenoptera of the Cynipidae family, has spread globally, from its native range in China, to Asia (1940), USA (1970), and Europe (2000), causing significant damages and economic losses in chestnut orchards and natural forests [23–26]. In the last twenty years, it has broadly spread from Italy to most of Eastern and Western European countries, including France, where it was first reported in 2005 in the Cevennes area before it reached the Pyrenees in 2013 [27]. In a recent ecological and genomic field study performed in the French Eastern Pyrenees, we provided the first quantitative estimate of the high invasive potential of this non-native insect and shown that it had spread across the entire fragmented chestnut tree population located in this area [22, 28]. As acknowledged in another recent study of *D. kuriphilus* invasion led in Northern Spain, the effect of the suitability and spatial configuration of the chestnut tree habitat on the spread of the pest remains poorly understood [29].

To improve our understanding of those determinants at a local scale, we first identify habitat patches based on the distribution of chestnut trees using a community detection algorithm (OSLOM algorithm [30]). We then apply graph traversal algorithms on the identified network of habitat patches to model the spread of the insect pest, which allows us to define and compute two complementary scores characterizing susceptibility to invasion. The first score, called ‘dangerousness’, quantifies the potential for a patch to spread the infestation, establishing a ranking of patches in terms of threats. The second score, ‘vulnerability’, similarly ranks patches but measures their susceptibility to infestation, regardless of which patch is initially affected. These scores combined with graph theory tools and susceptibility of the environment enable information extraction without prior assumptions about the composition of the forest landscape. The projection of these scores onto a map of the invaded area makes it possible to identify the spatial distribution of ‘dangerous’ and ‘vulnerable’ areas, leading to the characterisation of regions of interest for management of the invasive *D. kuriphilus* in French Eastern Pyrenees.

## Methods

### Modelling the spatial distribution of forests resources

The study area is located in the French Eastern Pyrenees and covers around 1,554 km^2^ of chestnut tree forests. Data from the national forest inventory [31] shows that *C. sativa* is one of the most abundant local species, although its distribution is fragmented and heterogeneous. The highly contrasted landscapes spanning from the Mediterranean Sea to the Canigó Massif (2,784 m above sea level) are indeed made of diverse vegetal formations associated to various densities of chestnut trees (see Fig. 1). Using data available online from IGN (National Institute of Geographic and Forest Information) [31], we combined the various layers of vegetal formations to obtain a partition of the spatial distribution of the forest resource. A study of the National Forest Inventory [32] reports the number of trees and species composition for each parcel in the French Eastern Pyrenees. Using this document and an exhaustive survey of the tree population in the studied area, we obtained in Table 1 an estimation of the number of chestnut trees for each of the six vegetal formations [31]. Together with the map of Fig. 1, we have estimated the number of chestnut trees per quadrats of 1 km^2^.

**Table 1.**
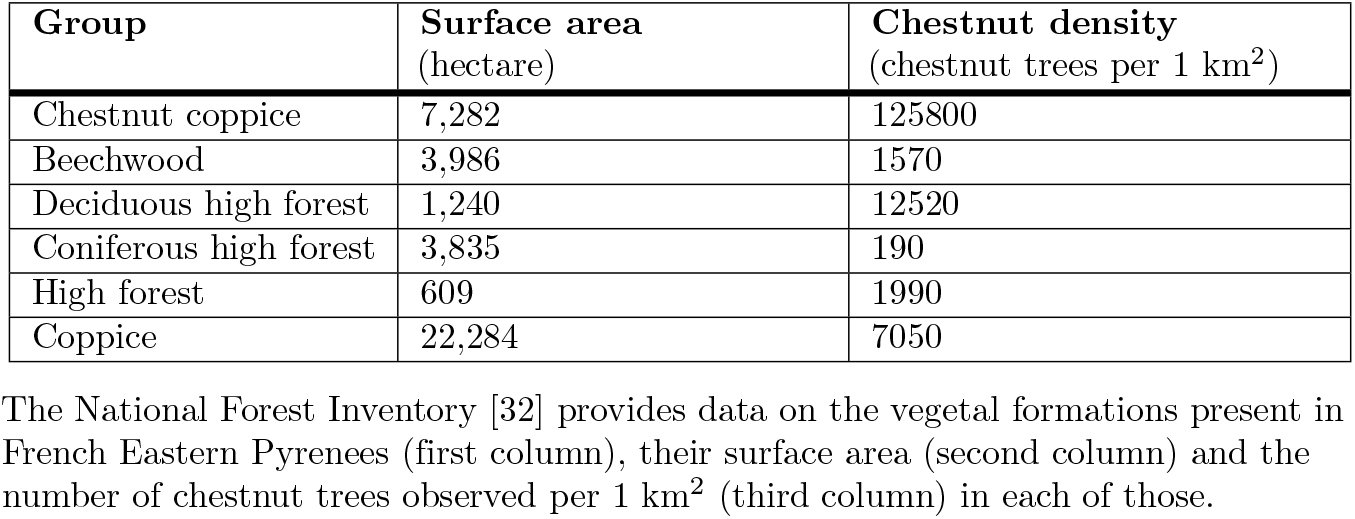
Estimation of the density of chestnut trees in French Eastern Pyrenees.

| Group | Surface area<br>(hectare) | Chestnut density<br>(chestnut trees per 1 km<sup>2</sup>) |
| --- | --- | --- |
| Chestnut coppice | 7,282 | 125800 |
| Beechwood | 3,986 | 1570 |
| Deciduous high forest | 1,240 | 12520 |
| Coniferous high forest | 3,835 | 190 |
| High forest | 609 | 1990 |
| Coppice | 22,284 | 7050 |

The National Forest Inventory [32] provides data on the vegetal formations present in French Eastern Pyrenees (first column), their surface area (second column) and the number of chestnut trees observed per 1 km<sup>2</sup> (third column) in each of those.

**Fig 1.**
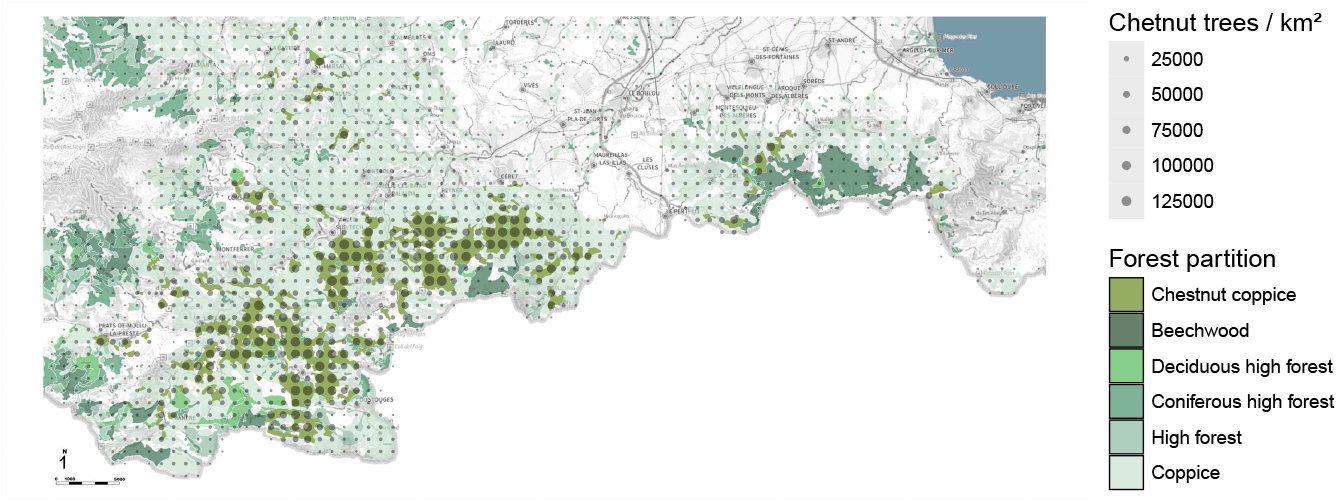
Spatial distribution of chestnut trees in French Eastern Pyrenees. Colours represent the different vegetal formations while the size of the dot indicates the density of chestnut trees per 1 km^2^ quadrat (see Table 1 for details). The south of the studied area is delimited by mountain peaks that defined the Spain border. The latter limits the study area because the forest resource is very different on the Spanish watersheds.

A *graph* is a mathematical object that represents relations among a set of entities. The entities are called *nodes*, and their relations are called *links*, typically depicted as lines between pairs of nodes. Here, the purpose of the graph is to represent spatial connections between the 1 km^2^ *quadrat* of forest resources. Accordingly, the nodes are the quadrats, and the links are the connections that are intended to model the propagation potential between two neighbouring quadrats. Some of these potentials are stronger than others, which is represented by a *weight* associated with every link.

With this representation, efficient propagation paths can be identified in the network of quadrats, with the aim of explaining and predicting how invaders are likely to spread through the forest. It is also possible to identify critical regions that should be monitored (or acted upon) in priority, based on various metrics from network analysis. Such tools have long been used in the study of social networks (see, e.g. [33]); they have also been used more recently in biological studies (see, e.g. [34]). In this study, we are particularly interested in the detection of so-called *communities*, which are clusters of nodes that are tightly connected to each other. In principle, such communities would correspond to patches in which the propagation potential between nodes is similar.

The structure of the graph itself is rather simple with a link between every pair of quadrats whose distance is less or equal to 3 km. The threshold of 3 km is a realistic assumption given the typical travel distances of the insect pest during their short adult lifetime, and as we do not intend to model the rare events of long-dispersion [29]. The most critical component in our modelling is the way weights are assigned to the links in order to represent the propagation potential of the invasive species depending on the features of both the parasite and the landscape. Since a complete description of the complex and intertwined behavioural and ecological mechanisms underlying such a potential remains elusive, we designed six alternative phenomenological metrics accounting for various hypotheses on the parasite flying capacity and on the suitability of the source and target habitats. Three of the metrics define weights that are identical in both directions, the other three can be asymmetrical. As a result, we considered both undirected graphs and directed graphs, depending on the underlying hypothesis, the directed cases have a different weight for both directions between a pair of neighbour nodes. The six metrics are listed in Table 2. They are all formulated in terms of three variables, namely *s, e*, and *d* defined as follows:

**Table 2.** Table of metrics associated to the edges of the graph.

| Metric | Edge weight | Graph |
| --- | --- | --- |
| 0 | $s \times e$ | undirected |
| 1 | $s \times e/d^2$ | undirected |
| 2 | $\sqrt{s} \times e$ and $\sqrt{e} \times s$ | directed |
| 3 | $\sqrt{s} \times e/d^2$ and $\sqrt{e} \times s/d^2$ | directed |
| 4 | $s$ and $e$ | directed |
| 5 | $s/d^2$ and $e/d^2$ | directed |

The metrics are computed according to the values of $s$ or $e$ that are the ratio of chestnut trees of the starting and ending node and $d$ the distance between the nodes.

- *s ∈* [0, 1] is the ratio of chestnut trees in the starting node of an edge
- *e ∈* [0, 1] is the ratio of chestnut trees in the ending node of an edge
- *d* is equal to the normalized Euclidean distance between nodes *s* and *e*, seeing the map as a grid where each quadrat defines an integer coordinate (quadrats sharing a side are at distance 1).

As the maximum number of chestnut trees found in a quadrat of 1 km^2^ is 125,800 (Table 1), we defined the density ratio of a quadrat as the number of chestnut tree over this number (e.g. a ratio of 0.5 corresponds to 62,900 chestnut trees par 1 km^2^).

The proposed metrics are meant to represent basic principles underlying the dispersal of invasive insect pests through a landscape where the host heterogeneity (and proximity) is described through *s, e*, and *d*. First, the potential number of *D. kuriphilus* individuals that emigrate from a starting quadrat could increase with the population of chestnut tree hosts in this quadrat (parameter *s*). Assuming that every individual parasite has a probability to leave that is independent of the host tree availability, the overall rate of propagation from a quadrat shall increase with the local abundance of parasites – which is indeed expected to rise linearly with *s* (metrics 0, 1, 4 and 5).

Alternatively, if the individual tendency to leave is lower in suitable habitats that provide host resources, the increase of the overall rate of propagation with the number of available hosts *s* is expected to become sublinear – which we model by considering the square root of *s* (metrics 2 and 3). Second, the introduction rate of invasive pests in a target quadrat is likely to increase with the number of chestnut tree hosts in this quadrat (parameter *e*) through habitat selection and, in the absence of interference between parasite individuals, such an increase is expected to be linear with *e* (metrics 0 to 3). Alternatively, in the case of passive dispersal [35], such an introduction rate may not depend on the suitability of the quadrat (metrics 4 and 5). Third, the rate of propagation between two quadrats decreases with the distance between them (parameter *d*). Since the surface of a two-dimensional area is proportional to the square of its diameter, the rate of propagation is considered to be inversely proportional to *d*^2^ (metrics 1, 3 and 5) (see also [36]). In order to single out the influence of the spatial isolation between quadrats, we also consider the ‘neutral’ assumption that the rate of pest propagation does not depend on *d* (metrics 0, 2 and 4). Finally, for each metric, we normalize all the resulting weights to values between 0 and 1.

### Community detection and supergraph

Community detection is a popular method for analysing the structure of a network. In the present case, these types of algorithms enable the identification of clusters of quadrats which we will refer to as patches, which are highly connected internally. There exist several definitions communities in the literature (see e.g. [37] for comparative study). The most classical definition is that a community is a group of nodes (a subgraph) that have more internal edges among them than they do collectively with the rest of the graph [38, 39]. Communities are also often defined algorithmically, in terms of the final products of some algorithm, without a structural definition [39]. However, all the definitions convey to some extent the fact that, within a community, the nodes are highly connected, which in our case means that a fast propagation of the pest can occur among the nodes of the same community. In other words, by analysing the community structure of the graph, we aim to capture patches (made of quadrats) in which the propagation is fast, while separating the patches between which the propagation is more difficult. To this aim, we applied a standard community algorithm called OSLOM [30] that clusters the map into communities, for each of the six graphs.

The patches identified by using the OSLOM community algorithm were then connected between each other to set a supergraph. In such a graph each node corresponds to a community, and every two nodes are connected according to the links between their respective quadrats. The weights on the edges were then defined as the inverse of the sum of the weights linking the quadrats belonging to the corresponding communities, so that small costs represent fast propagation. This higher scale of modelling enables the study of relationships between communities, particularly their vulnerability to cross-infection, with the aim of anticipating the spread of infection from one community to another. We used the shortest path algorithm developed by Dijkstra [40] to estimate how fast the propagation can occur between two remote communities and what are the most critical pathways in this propagation. Each node in the graph is at a certain distance from another node considered as the starting point. The Dijkstra algorithm permits to have a list of values of the shortest distances. These values are used to sort the other nodes of the supergraph (the patches). Complementary to the distance, the **betweenness centrality** is a measure of the importance of a node in a network based on its role as an intermediary in shortest paths. For instance, the betweenness centrality measures how a node is central in a graph by counting the number of the shortest paths connecting every two nodes that contains the central node. The greater the number of shortest paths through a node, the greater its centrality. Calculated for each node of the supergraph, this allows us to study the position of the nodes in the propagation pathways. Thus, the algorithm is the following one: for each community *c* of the supergraph, the Dijkstra algorithm is used to order all the other communities according to the cost of the shortest path to reach them from *c*. The rank of a community in this order can be seen as a priority criterion for watching certain patches of forests if *c* is infested. In addition to OSLOM program, calculations use `Python` [41], `ggraph` [42], `tidyverse` [43] and `NetworkX`[44] libraries.

### Dangerousness and Vulnerability

Following this calculation, six supergraphs are obtained; they detail the connections and pathways across the chestnut tree map of the French Eastern Pyrenees forest. We thus identified the most *dangerous* locations; namely, the locations that have the potential to quickly contaminate a considerable number of other locations. Conversely, we also identified locations that are most *vulnerable*; namely, the most likely to be reached quickly by several others in the event of a potential infestation. Computing results across the six supergraphs enables the analysis and comparison of the most dangerous and/or vulnerable locations for each metric.

- The **dangerousness score** of a community is defined as the average shortest-path cost from that community to all others. Communities are ranked accordingly, and the rank is normalized by the total number of communities, yielding a score between 0 and 1 across the six supergraphs. Each quadrat then inherits the score of its community (or the average score if it belongs to multiple communities).
- The **vulnerability score** of a community is defined as the normalized average rank (between 0 and 1) of its shortest-path distances from all other communities, reflecting how quickly it can be infested. Each quadrat inherits the score of its community (or the average if it belongs to multiple communities).

These scores are projected onto twelve maps, and a summary of these maps was constructed by counting the number of occurrences of each quadrat on the map according to two categories of values: score *<* 0.25 and ≤ 0.75 score. To have a global overview over the two scores (vulnerability and dangerousness), we computed an average value from the six graphs depending on the chosen propagation metric (Table 2). These average scores make it possible to highlight specific areas where the metrics generally agree on a critical level of dangerousness or vulnerability (score *>* 0.75), and similarly for areas where the metrics agree on a low risk (score *<* 0.25). A map per metric is also provided, so that these conclusions can be refined for a particular propagation model.

We defined **superspreaders** quadrats using the distance from the origin to a point with the two scores as coordinates. Thus, the quadrats are sorted using this distance, and we can extract list of top 5%, top 10% and top 20% quadrats with superspreading capabilities. We compared the dangerousness and vulnerability scores depending on the metrics with pairwise comparisons to evaluate the impact of the metric’s hypothesis on the results of the algorithms. We also compared the average dangerousness with the average vulnerability to emphasize the differences between both scores leading to a classification of quadrats that is projected to the map. This is complemented by other comparisons including the forest resource with the estimated number of chestnut trees in each quadrat that has been normalized to have values between 0 (no chestnut trees) and 1 (the maximum value). Consequently, average scores of dangerousness and vulnerability have been compared to the forest resource to highlight the impact of community clustering on the spread depending on density. These comparisons are done with the GGally package [45] allowing statistical and visual comparison of our normalized data.

## Results

### Community detection, supergraphs and patches

The modeling process produces six graphs (one per metric) that are made of 1,554 nodes with 8,998 edges (for undirected metrics) and 17,996 edges (for directed metrics). From each of the networks created according to the six metrics (Table 2), we obtained a supergraph using OSLOM community detection algorithm. Two example maps of the chestnut forest in the French Eastern Pyrenees are represented in Fig. 2. Some communities coincide in both maps (see e.g. communities 1, 37, 11 in the first map compared to 0, 33, 28 in the second map) and the number of communities also coincides mostly (see Table 3). However, this is not always the case (see e.g. in Fig. 2, communities 30 in the first map, versus 16 and 26 in the second). These divergences are an indicator that the choice of metric matters in the analysis.

**Table 3.** Number of communities detected with OSLOM algorithm.

| Metric | Graph | Number of communities |
| --- | --- | --- |
| 0 | undirected | 38 |
| 1 | undirected | 35 |
| 2 | directed | 35 |
| 3 | directed | 36 |
| 4 | directed | 39 |
| 5 | directed | 33 |

**Fig 2.**
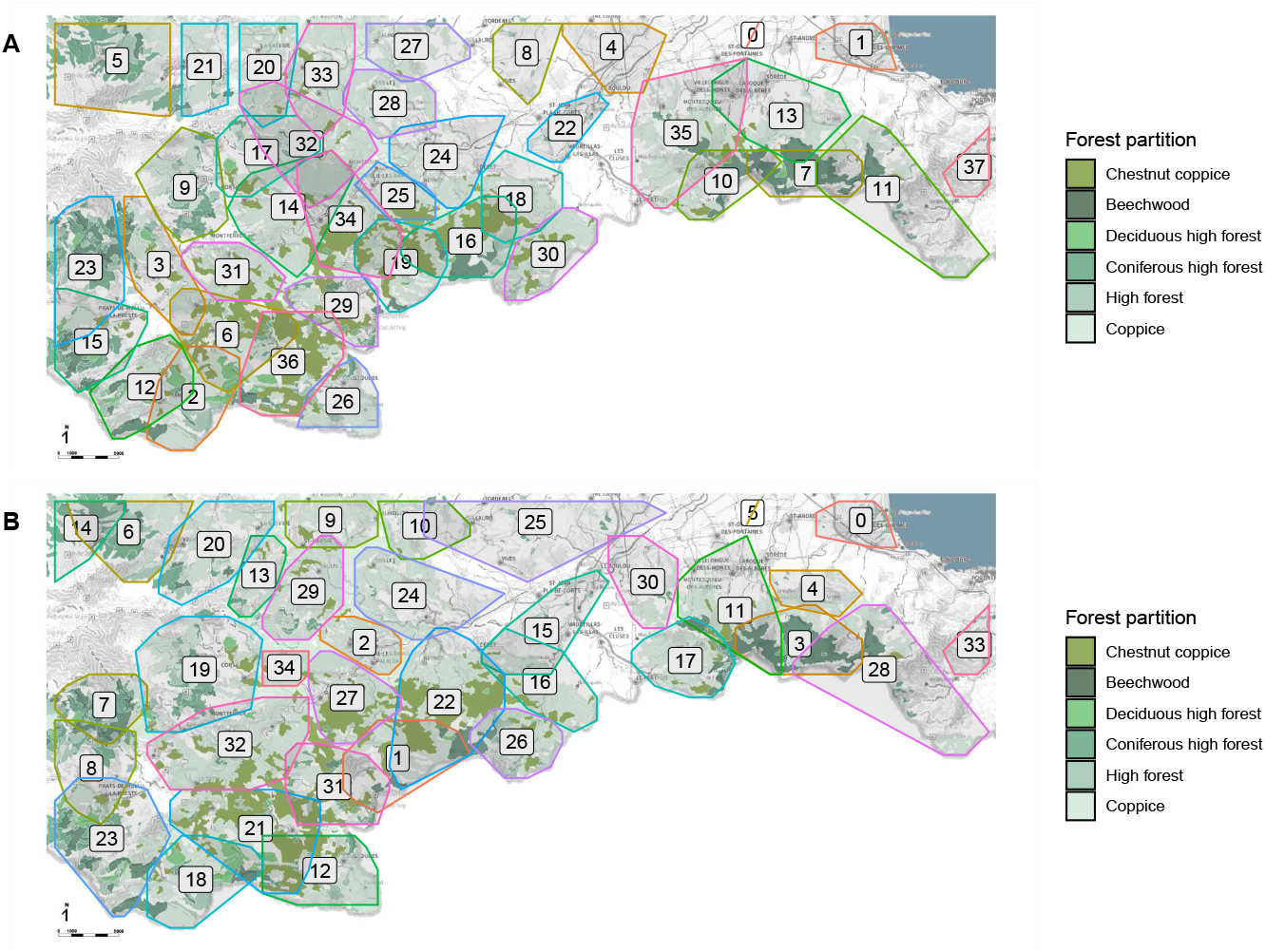
Community detection using OSLOM algorithm. The two examples of community detection presented here detect respectively 38 communities for the graph created using metric 0 (A) and 35 communities for the graph created using metric 1 (B).

For each possible starting node, the propagation through the graph of patches is estimated using the Dijkstra algorithm. For instance, Fig. 3 shows the community ranking from community 8, and even if communities 8 and 7 are close to each other, community 7 is not one of the first community because the value of the link between them is high.

**Fig 3.**
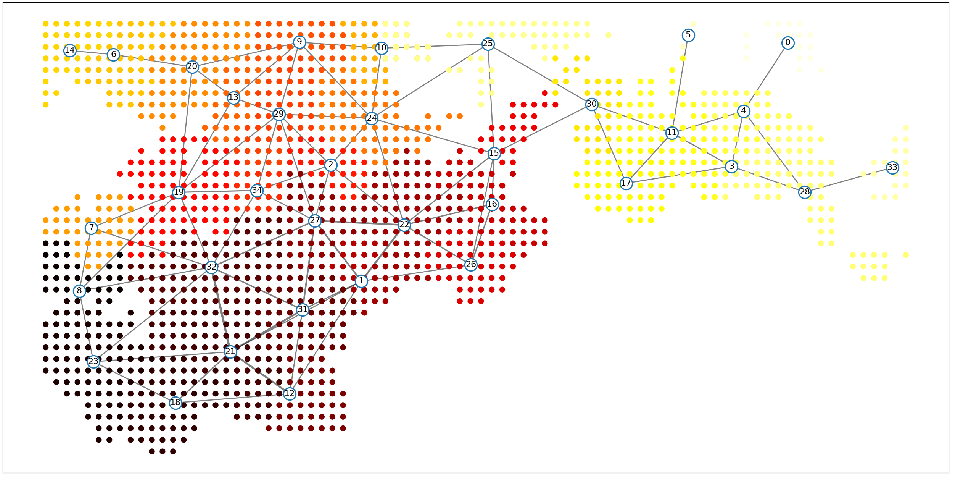
Supergraph over communities sorted with Dijkstra algorithm. Illustration of the communities ranking with community 8 as starting node for the invasion. If a node belongs to several communities, we compute the average rank to obtain the associated colour. In the red to yellow gradient for colours of nodes, dark red corresponds to the starting community and light yellow for the far one.

Basic statistics on these supergraphs (see supplementary material S1 Table for details) reveal that the overall structure is generally similar over all metrics. The main differences are observed on the supergraph of metric 4, where betweenness centrality shows a central node with a value above 500 and no other nodes with values between 400 and 500 (Fig. 4). This graph is also the only one showing a dense concentration of nodes and edges located in the top left side of the area. Supergraph 3 also exhibits some specificities, being the only one with a node of significant centrality (above 200) located in the upper-left area. This region appears to cover a large area, as there are no other nodes further north.

**Fig 4.**
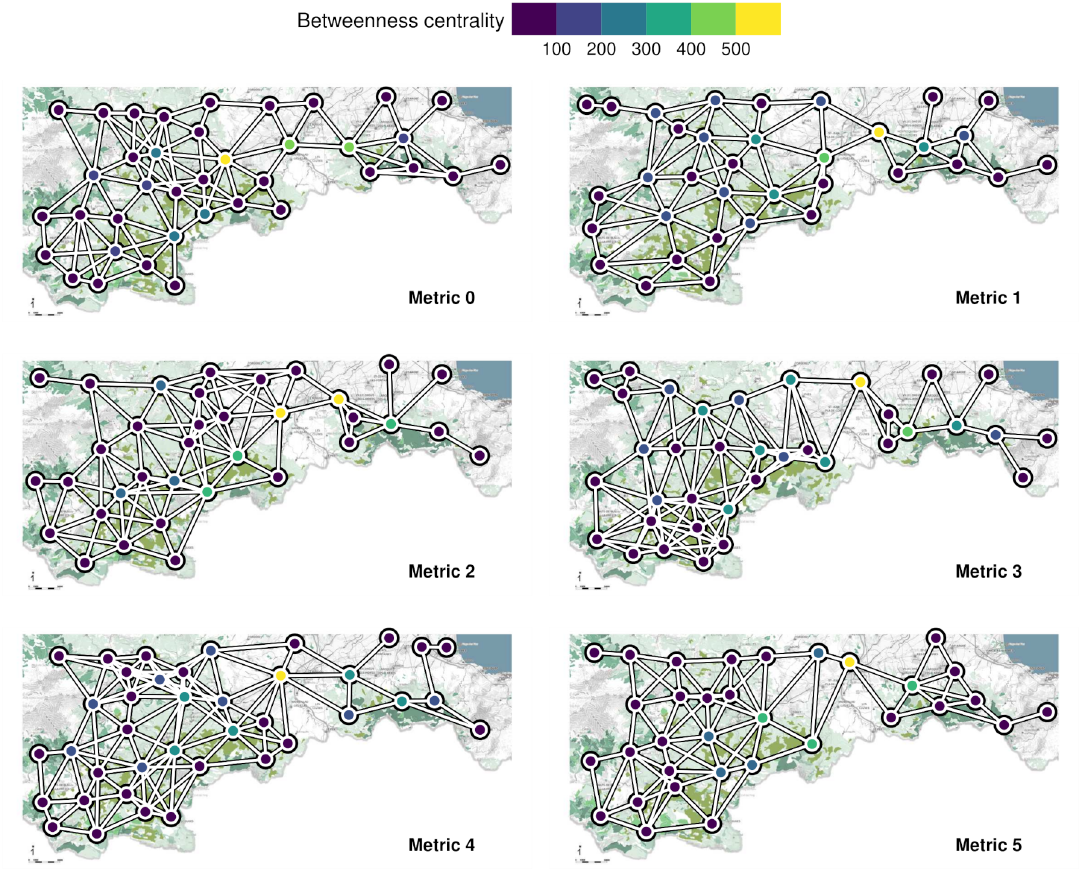
Betweenness centrality of the nodes of the six supergraphs. Edges of the supergraph are created if there are links between the communities in the graph.

#### Dangerousness and vulnerability maps

We made a pairwise comparison of the dangerousness scores of our six metrics including a normalized value of the number of chestnut trees for each patch. The scores established according to the different metrics are correlated with each other (with *r*^2^ values between 0.798 and 0.933) and, to a lesser extent, with the number of chestnut trees (*r*^2^ values between 0.486 and 0.553); details are available in supplementary material S1 Fig. Similar calculation on the vulnerability scores also indicates that the metrics are also correlated (with *r*^2^ values between 0.781 and 0.9) with also lower values with the number of chestnut trees (*r*^2^ values between 0.487 and 0.565); details are available in supplementary material S2 Fig. The six maps of dangerousness scores (supplementary material S3 Fig) and vulnerability (supplementary material S4 Fig) represent scores for the quadrats depending on the metric used ; Fig. 5 is a synthesis. Across most metrics (count *>* 3), 18.7% of quadrats record a dangerousness score below 0.25, compared with only 15.9% showing a vulnerability score below 0.25. Similarly (count *>* 3), 28.7% of quadrats correspond to the highest dangerousness class, compared with 26.1% classified as vulnerable. Of all the quadrats that are both very dangerous and very vulnerable, those that are both represent 87.7%. Despite few differences, metrics enable robust mapping.

**Fig 5.**
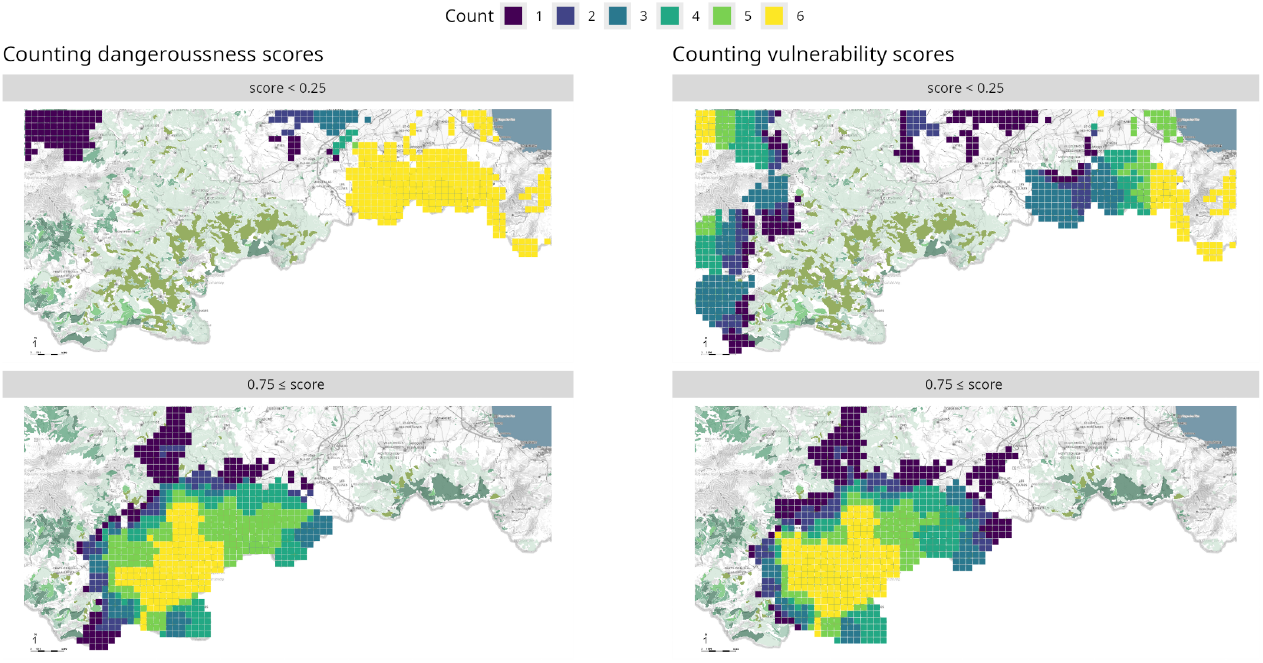
Map projection of categories of dangerousness and vulnerability scores. For each score (dangerousness and vulnerability) and each metric a partition in categories is made: the count is projected onto the map.

#### Comparison of vulnerability and dangerousness

The average vulnerability and dangerousness for each quadrat is calculated to assess the relationship between these two measures. Fig. 6A shows a strong correlation when dangerousness and vulnerability are higher than 0.5. Regarding values below the 0.5 threshold, it also distinguishes two areas where quadrats are more vulnerable than dangerous and reciprocally. This partition is projected onto the map (Fig. 6B) to delineate distinct areas within the study region. The east part of the studied forest is less dangerous than vulnerable, despite the presence of areas with a high concentration of chestnut trees. The area where dangerousness and vulnerability scores are high are located in the neighbourhood of the main chestnut forest in the south part of the studied forest. Superspreaders quadrats are identified because they are both dangerous and vulnerable (Fig. 6C). Beyond their spatial location, the map also conveys quadrat density, which underlies the partial correlation between chestnut tree resources and the scores.

**Fig 6.**
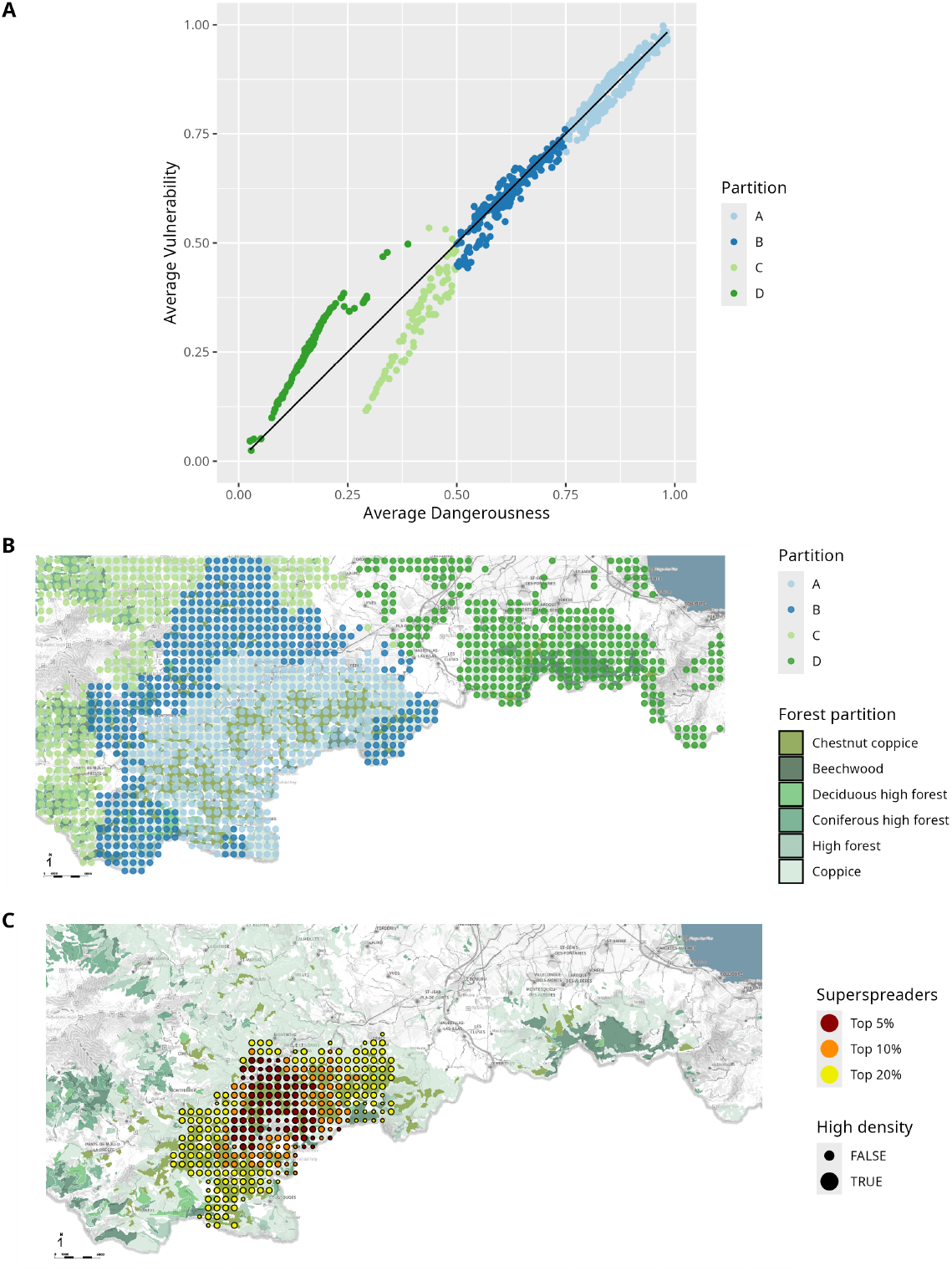
Correlation between average dangerousness and average vulnerability. Average dangerousness and vulnerability are computed from the six graphs depending on the metric (Table 2). A: Correlation of average values of metrics between dangerousness and vulnerability scores for each quadrat with a partition. B: Spatial distribution of the partition made on A to highlight dangerous quadrat without being vulnerable and vice versa. C: Superspreaders with high vulnerability and dangerousness. The density criterion is based on median value of chestnut tree resource (4132 chestnut trees per 1 km^2^).

## Discussion

We developed a tool to analyse the spatial distribution of chestnut trees and formulate predictions on the pest invasion pathways. This methodology is based on community detection to emphasize patches of the forest that share common properties.

Consequently, such patches experiment rapid colonisation by insect invaders, whose emigration to others forest patches is delayed. Further to this detection, we propose two scores derived on graph theory properties: dangerousness and vulnerability.

Dangerousness is a key component to underline contamination through communities because, by definition, it indicates that this community is often part of the shortest path that connects patches. Fig. 5 illustrates that the east forest of the map is not dangerous for the other because a very low scores are assigned to this forest. This is due to fewer connections between the communities; thus, it reduces the risk. This indicates that the propagation will mainly transit to this forest with the higher concentration of transmission pathways. These pathways are not identified with the betweenness measurements (Fig. 4) because of the topology of the network where the east-west transitions counter-balance the process in the east side preventing such identification. Maps per metrics also indicates some specificities (supplementary material S3 Fig). Especially, for metric 3, a corridor from north to south has been emphasized indicating that the surveillance of the north of the studied area could be relevant to prevent a transmission to the most densely populated chestnut forest in French Eastern Pyrenees. In addition, this corridor is closed to a natural barrier in terms of altitude because the presence of Canigó Massif preventing connection from the north to the south part of the map because this altitude lies outside the natural range of chestnut tree distribution.

The overview of vulnerability scores (secund column of Fig. 5) indicates the susceptibility of each quadrat to become infected, wherever the invasion starts. The less vulnerable areas are located at the extreme east and west sides, but the less dangerous are only located on the east side of the studied area. This partition is materialised by the valley by the highway from France to Spain that separates the two forest massifs. This result is coherent to the definition because all patches are considered as the starting point of the infestation and the score is an average ranking in terms of position in the transmission chain.

Fig. 6 yield a smoothing of the results while enabling a broader perspective, as an average value was computed for both scores. This approach with several possibilities of metrics avoids the pitfalls that raw data can generate by training the algorithms towards singular solutions. The comparison between the two average scores (vulnerability and dangerousness) is very relevant (Fig. 6) because it underlines some particular aspects of the geographical distribution of the chestnut trees resource over this territory. Such a classification is reported to the map (Fig. 6B) underlying the possible invasion scenario from the north to the south belonging to group B where both values of dangerousness and vulnerability are between 0.5 and 0.75. For a given set of 1 km^2^ quadrats, vulnerability is strongly correlated to dangerousness leading us to the identification of superspreaders (Fig. 6C). However, this pattern does not hold for peripheral chestnut trees, highlighting areas to prioritize for protection versus those requiring less monitoring. Consequently, this map combined with the community map (Fig. 2) could lead to define specific area in the forest to monitor invasion progression and quantify the infestation due to their vulnerability because of these specificities. The least dense quadrats within the identified superspreaders have been detected (Fig. 6C), they will quickly become infested but will be easier to monitor. This specific distribution could lead to a delimitation that could serve to evaluate the risk of invasion and give insight for preserving these vulnerable quadrats. We thus propose to monitor actively the least dense quadrats to have an early detection of a potential infestation.

While statistical relationships have been shown between infestation levels and the frequency of chestnut trees in the population [22, 46], our hypothesis states that quadrats with few chestnut trees within highly vulnerable communities should be infested as quickly as quadrats with a high number of chestnut trees. As a low abundance of chestnut trees is easily monitored, these quadrats show a promising potential for the early detection of cynips galls. Our theoretical results obtained from the map are consistent with the geographic surveys conducted in these forests in recent years [22] where the infestation has been quantified.

Leveraging the explicit spatial distribution of chestnut trees with our metrics can be seen as a complementary approach to aid in establishing biological control with *Torymus sinensis* [27]. Contrary to dynamical theoretical models using partial differential equation describing two types of populations (*D. kuriphilus* and *T. sinensis*) in [47] or a short and long range dispersal model through Europe in [48], our methodology provides maps with scores, counts and superspreaders. Our results contain additional knowledge computed from spatial distribution of trees that might enrich the dynamical models previously mentioned. With a similar approach using graph to represent connection between patches, Perry *et al*. [21] proposed to select control location using betweenness centrality. In our case, their procedure was not applicable to our graph with a low clustering coefficient compared their structure that is highly clustered leading to clear control points in the graph (supplementary material S1 Table and Fig. 4 for details on supergraphs structures).

One of the limitations of our approach is its computational cost, which depends on the size of the graph. For example, the time complexity of community detection in a graph of n nodes and m edges using the Girvan-Newman algorithm [49] is proportional to *nm*^2^. As the size of the graphs increase, such algorithms become rapidly inefficient. In comparison, the OSLOM algorithm has a running time proportional to *n*^2^, which is faster. For a recent review of community detection methods with computational complexity analysis see [37]. Note that the metric choices previously made ensure a reasonable running time for the problem addressed here. Depending on their computational resources, the users of this approach may set a different spatial resolution of the quadrats to obtain a network of manageable size at an affordable computational cost.

## Conclusion

We provide a tool analysing the spatial distribution of a specific forest tree (*C. sativa*) over a territory. The objective is to measure encountered risk of a parasite infestation (*D. kuriphilus*) to this forest resource with agronomical interest. From the map, we built a graph connecting a set of quadrats (1 km^2^) partitioning the forest, and we define six metrics to have various estimates of the colonisation pathways of the parasite through the network of chestnut trees. A community detection algorithm splits the forest in patches depending on a set of parameters, leading to six partition maps. These maps provide details on forest areas that should be considered strongly connected because they belong to the same community. Further to these partitions, we provide a set of two scores that quantify the dangerousness of a quadrat and its vulnerability to parasite colonisation. Projected onto a map, the quadrats with high and low scores could be considered as helpers to define a cartography of the area to protect to anticipate and react to parasitic infestation. This topological interpretation of pathways can also be interpreted as scenario by the decision-makers and forest park managers to help them to define strategies. Our main result is that we can establish maps of dangerousness and vulnerability for an invasive insect using the spatial distribution of its tree resource leading to the authentication of superspreaders. These maps predict potential pathways of invasion and can be used to improve the early detection and the monitoring of an invasion. These maps are not strongly correlated to the distribution of forest resources, bringing a betterment regarding data available from maps. Finally, we propose to monitor specific areas with few chestnut trees that are part of superspreaders to facilitate the detection of the early stages of an invasion. The adaptation of this methodology to well-mapped and particularly vulnerable agro-ecosystems offers promising potential for crop pest management.

## Supporting information

**S1 Table.** Statistics of the six supergraphs.

| Metric | Order | Size | Diameter | Efficiency | Average distance |
| --- | --- | --- | --- | --- | --- |
| 0 | 38 | 224 | 9 | 0.40 | 3.54 |
| 1 | 35 | 185 | 10 | 0.39 | 3.72 |
| 2 | 35 | 216 | 9 | 0.42 | 3.43 |
| 3 | 36 | 226 | 10 | 0.40 | 3.72 |
| 4 | 39 | 256 | 10 | 0.42 | 3.39 |
| 5 | 33 | 194 | 10 | 0.41 | 3.63 |

These metrics are specific measures or properties of the graph, used to compare or analyse its structure:

- Order: the number of vertices in the graph.
- Size: the number of edges in the graph.
- Diameter: the greatest distance (shortest path length) between any two vertices.
- Efficiency: a measure of how efficiently the graph transmits information, often defined as the average of the inverses of the shortest path lengths between all pairs of vertices.
- Average distance: the mean shortest path length between all pairs of vertices.

**S1 Fig.**
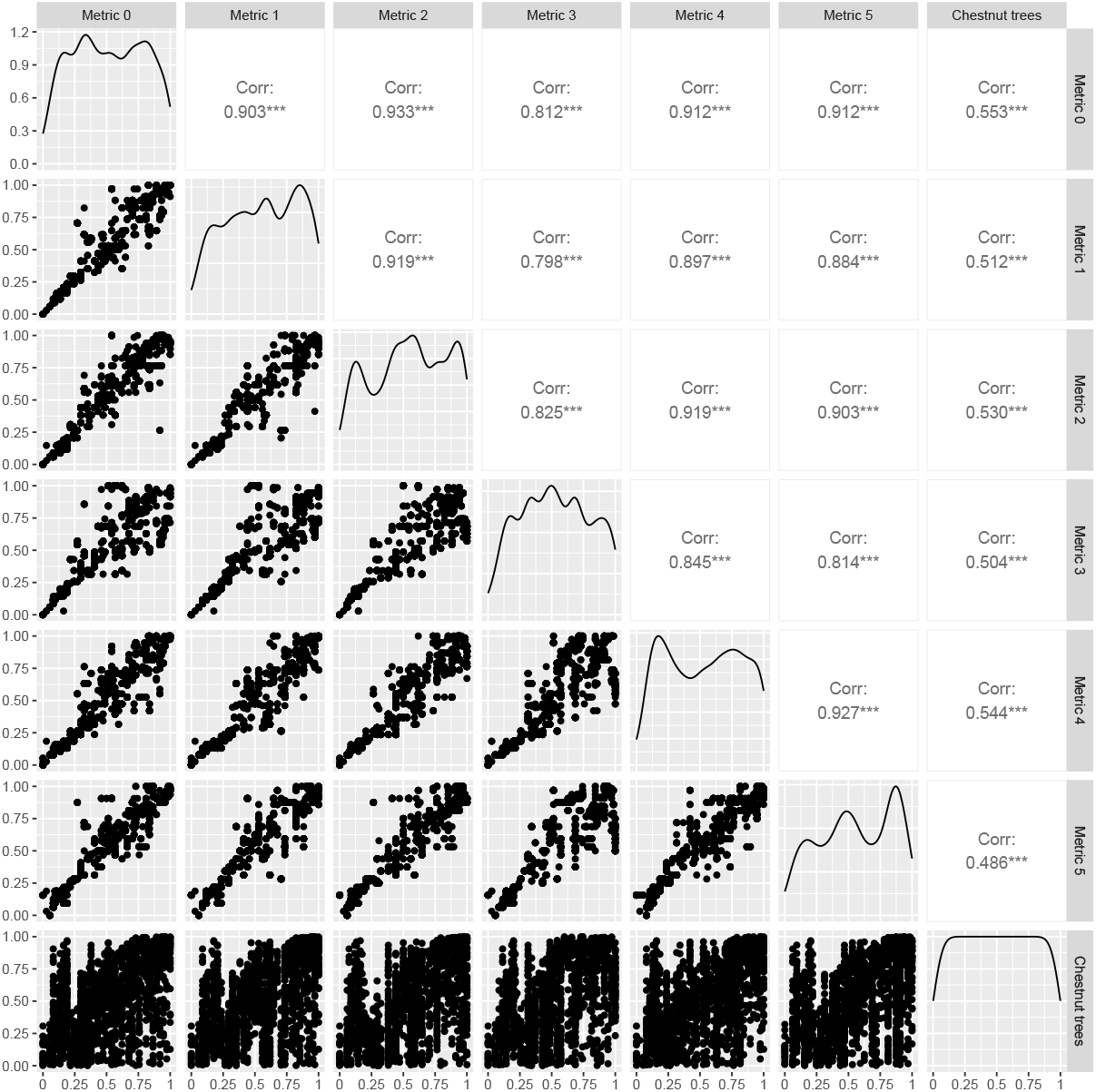
Pairwise comparison and correlation between dangerousness scores and forest resource of quadrats. It represents the correlations (visual and statistical) between metrics and a normalized value of the number of chestnut trees in a quadrat. All the scores are correlated: the dangerousness for the six supergraphs and the forest resource. The correlations of metric 3 are below the others with an average value of 0.812 while the other correlations are closer or higher than 0.9. This correlation is statistically valid for the number of chestnut trees even if the correlation value is close to 0.5.

**S2 Fig.**
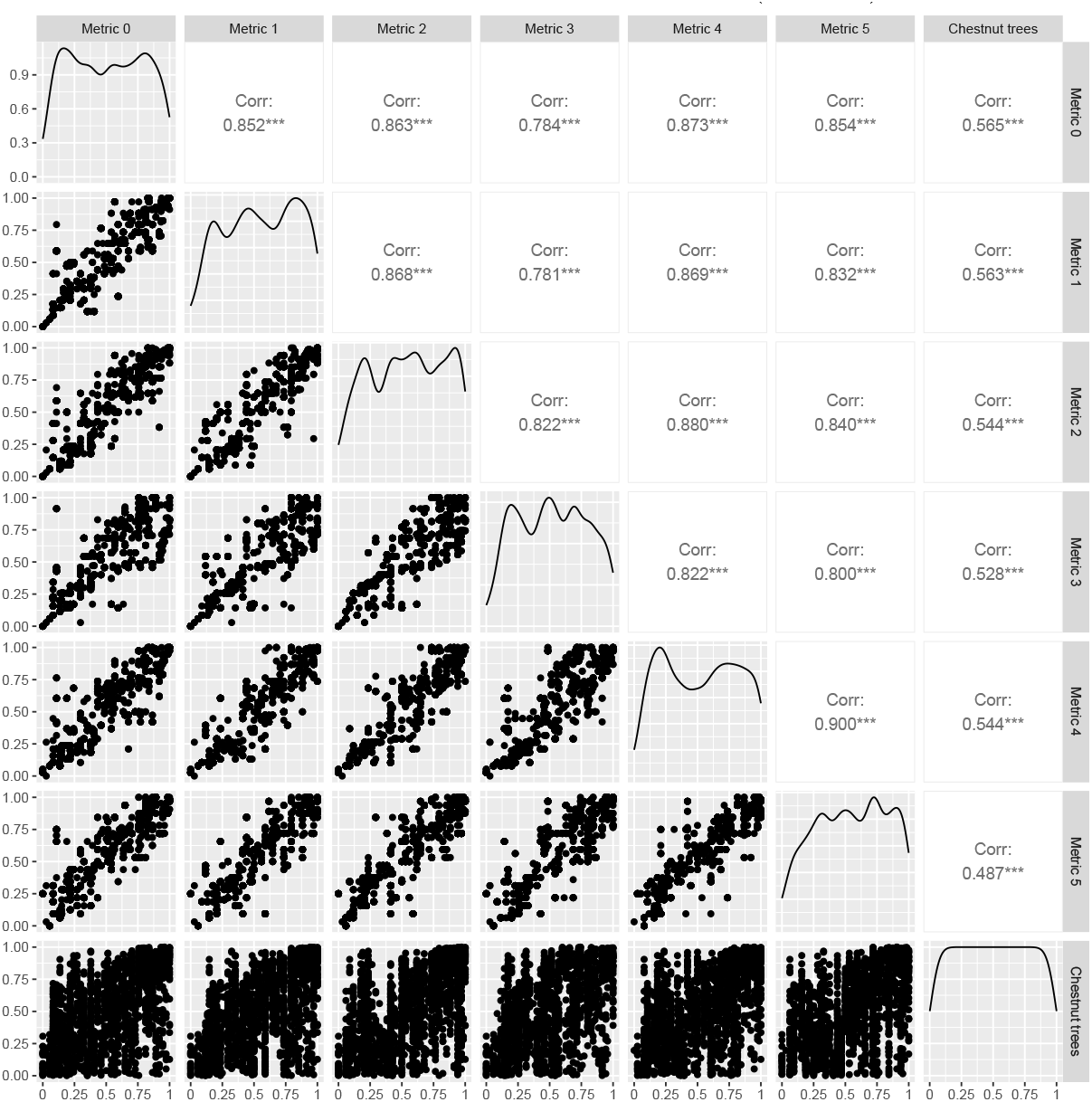
Pairwise comparison and correlation between vulnerability scores and forest resource of quadrats. It represents the correlations (visual and statistical) between metrics and a normalized value of the number of chestnut trees in a quadrat. Metrics 4 and 5 are correlated with a value of 0.927 but, regarding the correlation of these both metrics with the ranked chestnut trees, the values are equal to 0.54 and 0.48, respectively. This underlines that the number of chestnut trees really impacts the scores based on the definition of the two graphs where the metric 4 depends only on the source resource and where the distance is included for metric 5 (Table 2).

**S3 Fig.**
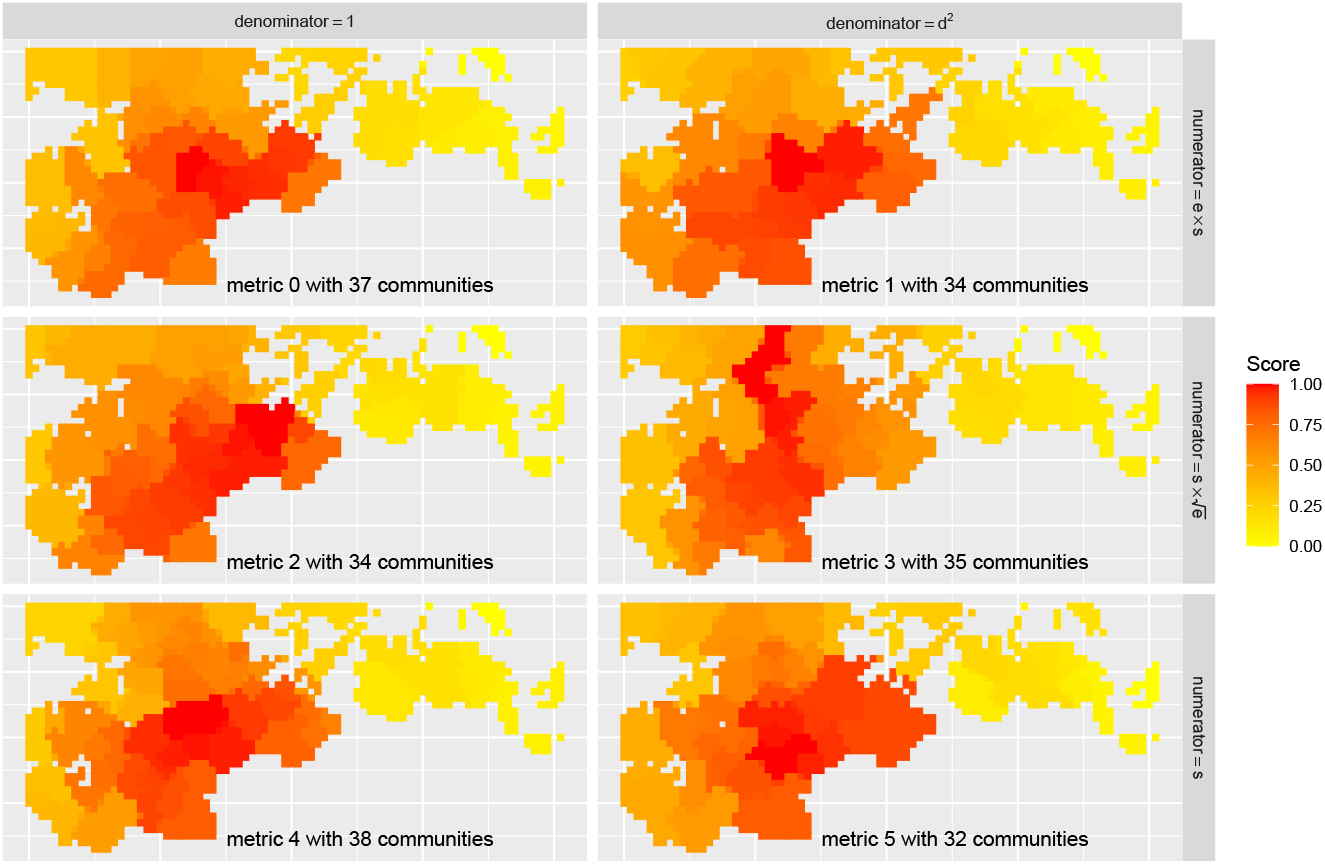
Projection of dangerousness scores for each metric. Dangerousness scores are computed for the six graphs depending on the metric (Table 2) where lines are for the numerator and columns for the denominator of the metric. Dangerousness scores of each quadrat are represented with the colour gradient (from yellow to red) shown in the legend. Except for the metric 3, the most dangerous area is consistently located in the south centre of the studied chestnut tree population.

**S4 Fig.**
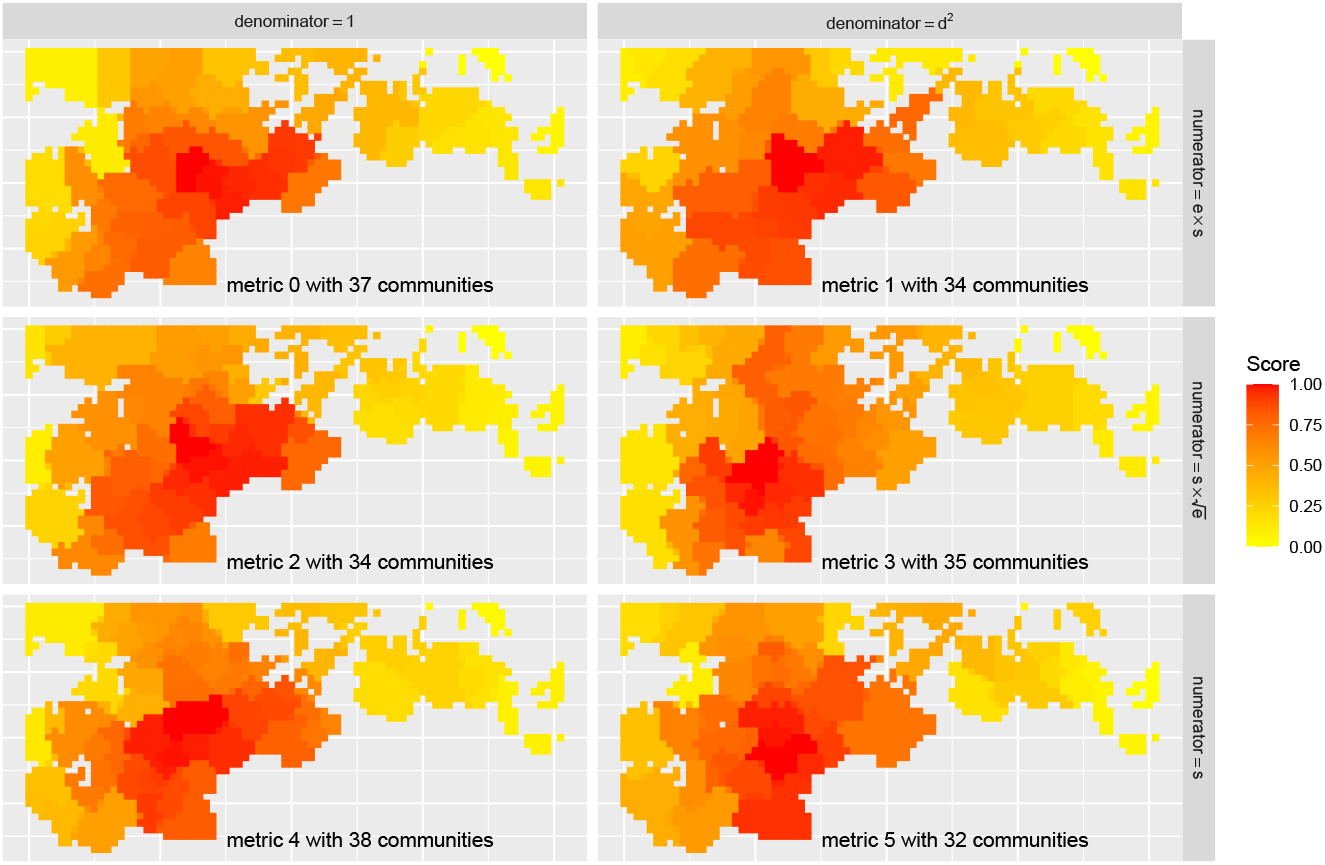
Projection of vulnerability scores for each metric. Vulnerability scores are computed for the six graphs depending on the metric (Table 2) where lines are for the numerator and columns for the denominator of the metric. Vulnerability scores of each quadrat are represented with the colour gradient (from yellow to red) shown in the legend.

## Acknowledgments

This study was supported by the MITI project: “Invasions biologiques et ressources hétérogènes : de la modélisation hybride á la gestion Eco-Bio-Sociale” founded by the Mission pour les Initiatives Transverse et Interdisciplinaires of the CNRS. This study is set within the framework of the “Laboratoires d’Excellences (LABEX)” TULIP (ANR-10-LABX-41) and of the “École Universitaire de Recherche (EUR)” TULIP-GS (ANR-18-EURE-0019).

## Author contributions

Conceptualisation: MR, AC, SG. Methodology: MR, AC, SG. Investigation: All authors. Visualisation: MR. Funding acquisition: MR, AC, SG. Supervision: MR, SG. Writing-original draft: MR. Writing-review & editing: All authors.

## Data and materials availability

All data needed to evaluate the conclusions in the paper are present in the main text and/or the Supplementary Materials.

